# Apolipoprotein L1 Dynamics in Human Parietal Epithelial Cell Molecular Phenotype Kinetics

**DOI:** 10.1101/259267

**Authors:** Vinod Kumar, Himanshu Vashistha, Xiqian Lan, Nirupama Chandel, Kamesh Ayasolla, Shadafarin Marashi Shoshtari, Rukhsana Aslam, Nitpriya Paliwal, Frank Abbruscato, Joanna Mikulak, Waldemar Popik, Ashwani Malhotra, Catherine Meyer-Schwesinger, Karl Skorecki, Pravin C Singhal

## Abstract

Human Parietal Epithelial cells (PECs) are considered as a source of progenitor cells to sustain podocyte (PD) homeostasis. We hypothesized that the absence of apolipoprotein (APO) L1 favors the PEC phenotype and that induction of APOL1 transitions to PD renewal. During PECs’ transition, APOL1 expression coincided with the expression of PD markers (PEC transition) along with down regulation of miR193a. The induction of APOL1 down regulated miR193a and induced PD markers in PECs/HEKs; whereas, the APOL1-silencing in transited (Tr)-PECs/HepG2s up regulated miR193a expression suggesting a reciprocally linked feedback loop relationship between APOL1 and miR193a. HIV, IFN-y, and vitamin D receptor agonist (VDA) induced APOL1 expression and PEC transition markers but down regulated miR193a in PECs/HEKs. Glomeruli in HIV patients and HIV: APOL1 transgenic mice displayed foci of PECs expressing synaptopodin, a PEC transition marker. Since APOL1 silencing in PECs partially attenuated HIV-, VDA-, and IFN-y-induced PECs transition, this would suggest that APOL1 is an important functional constituent of APOL1-miR193a axis.

---

APOL1 (G0) is a minor component of circulating lipid-rich trypanolytic multiprotein complexes in certain primate species including humans (1). It is expressed in liver, pancreas, kidney, brain, macrophages, and endothelial cells (1). Approximately 34% of African Americans carry one of the two risk variants (G1 and G2) and 13% have both variants (2, 3). Increased expression of G1 and G2 risk variant APOL1, has been shown to induce cellular injury, including podocytes (PDs), both *in vitro* and *in vivo* (4–13).

Parietal epithelial cells (PECs) and PDs are derived from the same mesenchymal cells during embryogenesis (14–16); however, the expression level of miR193a determines the net phenotype- PECs vs. PDs- in these cells (17). We have recently demonstrated that APOL1-miR193a axis preserves podocyte molecular phenotype in adverse milieus (18). We now hypothesize that APOL1 and miR193a forms a reciprocally linked feedback loop to regulate PECs' phenotype in humans. In this loop, lack of APOL1 assures PECs’ phenotype while its presence initiates their transition to PDs.

## APOL1 and miR193a are reciprocally linked with a feedback loop in PECs

PECs have not been shown to display expression of APOL1 both *in vitro* and *in vivo* studies (1, 19). To examine the expression of APOL1, lysates of cultured PECs, differentiated PDs and HepG2 (positive control) and HEKs (negative control) were probed for APOL1 and GAPDH. HepG2s and PDs displayed robust expression of APOL1, but PECs and HEKs did not display any expression of APOL1 protein (Fig. 1A).

**Fig.1.**
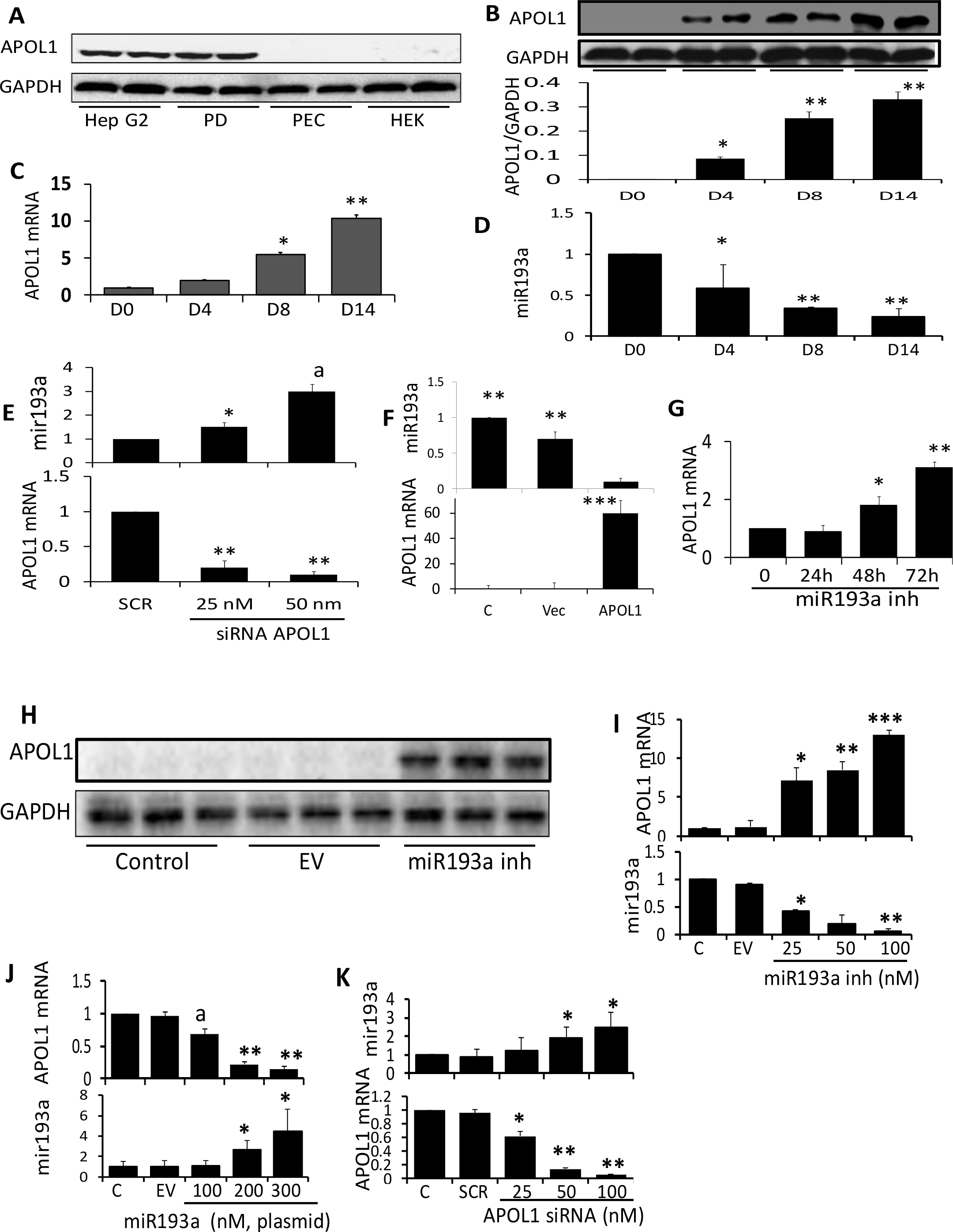
APOL1-miR193a Axis in Human PECs. A. Protein blots of PECs, HEKs, differentiated podocytes (PD), and HepG2s were probed for APOL1 and reprobed for GAPDH (n =3). Gels from two different lysates are displayed. PECs and HEKs did not show any expression of APOL1. On the other hand, both PDs and HepG2s displayed robust expression of APOL1. B. PECs were incubated in media for different time period (0, 4, 8, and 14 days [D]) at 37°C (n=4). Subsequently, protein blots were probed for APOL1 and reprobed for GAPDH. Blots from two different lysates are displayed. The lower panel shows cumulative densitometric data in bar graphs. APOL1 expression emerged on day 4 and progressed with time. *P<0.05 compared with D0; **P<0.01 compared with D0, and D4. C. RNAs were extracted from cell lysates of the experiments carried out in B (n=4). cDNAs were amplified with a specific primer to *APOL1*. Cumulative data are shown in a bar diagram. *P<0.05 compared to D0 and D4; **P<0.01 compared to D0, D4, and D8. D. MicroRNA193a was measured from RNAs extracted in C. Cumulative data are shown in a bar diagram. *P<0.05 compared with D0; **P<0.01 compared with D0. E. To determine the role of APOL1 on PECs’ miR193a expression, PECs were transfected with either scrambled (SCR) or miR193a-siRNA (25 nM or 50 nM) (n=3). RNAs were extracted. cDNAs were amplified with a primer specific for APOL1; cumulative data are shown in a bar diagram (Lower panel). The same RNAs were assayed for miR193a. Cumulative data are shown in the upper panel. *P<0.05 compared with respective SCR; **P<0.01 compared with respective SCR; ^a^P<0.01 compared with respective SCR and 25 nM. F. To confirm the role of APOL1 in regulation of miR193a expression, vector and APOL1 lentivirus was transduced in PECs (n=3). RNAs were extracted from control ( C), vector (Vec) and APOL1-tranduced PECs. cDNAs were amplified with a primer specific for APOL1. Cumulative data are shown in a bar diagram in the lower panel. miR193a was assayed from the extracted RNAs. Cumulative data are shown in a bar diagram in the upper panel. **P<0.01 compared with APOL1; ***P<0.001 compared with C and Vec. G. To evaluate the effect of miR193a on APOL1 expression, PECs were incubated in media containing a specific miR193a inhibitor (25 nM) for different time periods (0, 24, 48, 72 hours) (n=3). cDNAs were amplified with a primer specific for APOL1. Cumulative data are shown in a bar diagram. *<0.05 compared with 0 and 24h; **<0.01 compared with 48h. H. To confirm the effect of miR193a inhibitor on the induction of APOL1 protein, PECs were transfected with either empty vector (EV) or miR193a inhibitor (inh) (n=3). Proteins were extracted from control, EV- and miR193 inh-transfected PECs. Protein blots were probed for APOL1 and reprobed for GAPDH. Blots of three different lysates of control and experimental PECs are shown. I. To determine the dose response effect of miR193a inhibitor, PECs were transfected with either empty vector (EV) or different concentrations of miR193a inhibitor (plasmid, 25, 50, and 100 nM) (n=3). RNAs were extracted and assayed for miR193a. Cumulative data are shown in a bar diagram in the lower panel. cDNAs were assayed with a primer specific for APOL1. Cumulative data are shown in a bar diagram in the upper panel. *P<0.05 compared with respective C, SCR. **P<0.01 compared with respective C, EV, mir193a inh, 25 nM. ***P<0.001 compared with respective C and EV. J. To confirm the effect of miR193a, PECs were transfected with empty vector (EV) or different concentrations of miR193a plasmid (100, 200, and 300 nM) (n=3). RNAs were extracted and assayed for miR193a from control (C) and experimental cells. Cumulative data are shown in a bar diagram in the lower panel. cDNAs were amplified with a primer specific for APOL1. Cumulative data are shown in a bar diagram in the upper panel. *P<0.05 compared to respective C, EV, and miR193a, 100 nM; **P<0.01 compared with respective C, EV, and miR193a, 100 nM; ^a^P<0.05 compared to respective C and EV. K. To determine the regulatory role of APOL1 on miR193a in non-kidney cells, HepG2s were transfected with either scrambled (SCR) at different concentrations (25, 50 and100 nM) of APOL1siRNA (n=3). RNAs were extracted from control and experimental cells. cDNAs were amplified with a primer specific for APOL1. Cumulative data are shown in a bar diagram in the lower panel. RNAs were assayed for miR193a and cumulative data are shown in a bar diagram in the upper panel. *P<0.05 compared with respective C and SCR; **P<0.01 compared with respective C, SCR, and siRNA APOL1, 25 nM.

Since APOL1 is expressed by PDs (1, 4, 19), we asked whether PECs transiting to PDs’ phenotype would also display APOL1. PECs were incubated in media for different time intervals followed by RNA and protein extractions. Protein blots were probed for APOL1 and GAPDH. PECs did not show expression of APOL1 protein on day 0, however, APOL1 expression emerged on day 4 and increased further during transition (Fig.1B). RNAs were assayed for miR193a and cDNAs were amplified with an *APOL1* specific primer. *APOL1* mRNA expression increased (Fig. 1C) but miR193a levels decreased (Fig. 1D) during transition.

To evaluate a possible causal relationship, PECs were transfected with either scrambled or APOL1 siRNAs. cDNAs were amplified for *APOL1* and RNAs were quantified for miR193. Down regulation of *APOL1* mRNA (Lower panel, Fig. 1E) was associated with an up regulation of miR193a (Upper panel, Fig. 1E). These results suggest that PECs have a functional APOL1 protein which is not detectable by available tools; nonetheless, when knocked out up regulates miR193a expression.

For example a possible inverse relationship, PECs were transduced with vector or APOL1 (lentivirus). Protein and RNAs were extracted from Control (C), vector (Vec) and APOL1-transduced PECs. cDNAs were amplified for *APOL1* and RNAs were quantified for miR193a. An enhanced *APOL1* expression (Lower panel) was associated with down regulation of miR193a expression (Upper panel) (Fig.1F). APOL1-transduced PECs displayed enhanced APOL1 protein expression (data not shown).

If miR193a is also negatively regulating *APOL1* gene expression then down regulation of miR193a should also de-repress *APOL1* expression in PECs. miR193a expression in PECs was inhibited by a specific inhibitor and evaluated for *APOL1* mRNA expression. Inhibition of miR193a expression enhanced *APOL1* mRNA expression in PECs (Fig. 1G).

To confirm the effect of miR193a inhibitor on APOL1 protein expression, PECs were transfected with either empty vector or miR193a inhibitor (n=3). Protein blots were probed for APOL1 and reprobed for GAPDH. Blots of three different lysates of control and experimental PECs are shown in Fig. 1H. miR193a inhibitor induced APOL1 protein expression in PECs.

To assess a possible APOL1-miR193a axis in non-kidney cells, HepG2s were treated with a miR193a inhibitor, followed by extraction of RNAs and assayed for miR193a as well as *APOL1* mRNA. As shown in Fig. 1I, inhibition of miR193a in HepG2s (Lower Panel) resulted in an enhanced expression of *APOL1* mRNA (Upper Panel) in a concentration-dependent manner.

To confirm a reciprocal relationship between miR193a and APOL1 expression in HepG2s, cells were transfected with either empty vector (EV) or miR193a plasmids. cDNAs were amplified for *APOL1* and RNAs were assessed for miR193a. Enhanced miR193a expression in HepG2s (Lower panel) was associated with down regulation of *APOL1* mRNA expression (Upper panel, Fig. 1J). To validate feedback relationship, RNAs were extracted from control HepG2 and HepG2s-transfected with scrambled (SCR) siRNA or APOL1 siRNA. RNAs were evaluated for miR193a and *APOL1*. Down regulation of *APOL1* expression (Lower panel, Fig. 1K) was associated with up regulation of miR193a expression (Upper panel, Fig. 1K).

## Both absence and presence of APOL1 characterize PECs’ molecular profile

To characterize PECs’ molecular phenotype, proteins and RNAs were extracted from control (cultured at 33°C) and Tr-PECs (PECs seeded on fibrin-coated plates for 4-14 days at 37°C). Protein blots were probed for several PEC and PD markers and reprobed for GAPDH (n=3). Representative gels (on 0 and 14 days) are displayed in Figs. 2A and 2C. Densitometric data are shown in bar diagrams (Figs. 2B and 2D). PECs’ transition was characterized by down regulation of PEC markers, induction of APOL1, and an up regulation of transition markers. Representative fluoromicrographs displaying induction of APOL1 and transition marker are shown in Fig. 2E. cDNAs were amplified with specific primers for PECs and their transition markers. Cumulative data are shown in bar diagrams (Figs. 2F and 2G). To determine the time course effect on loss/gain of PECs/transition markers, PECs were incubated for different time periods at 37^°^C. cDNAs were amplified for PECs//transition markers and cumulative data are shown in Figs 2H and 2I. Tr-PECs displayed a decrease in PECs but several fold increase in their transition markers along with APOL1 expression.

**Fig.2.**
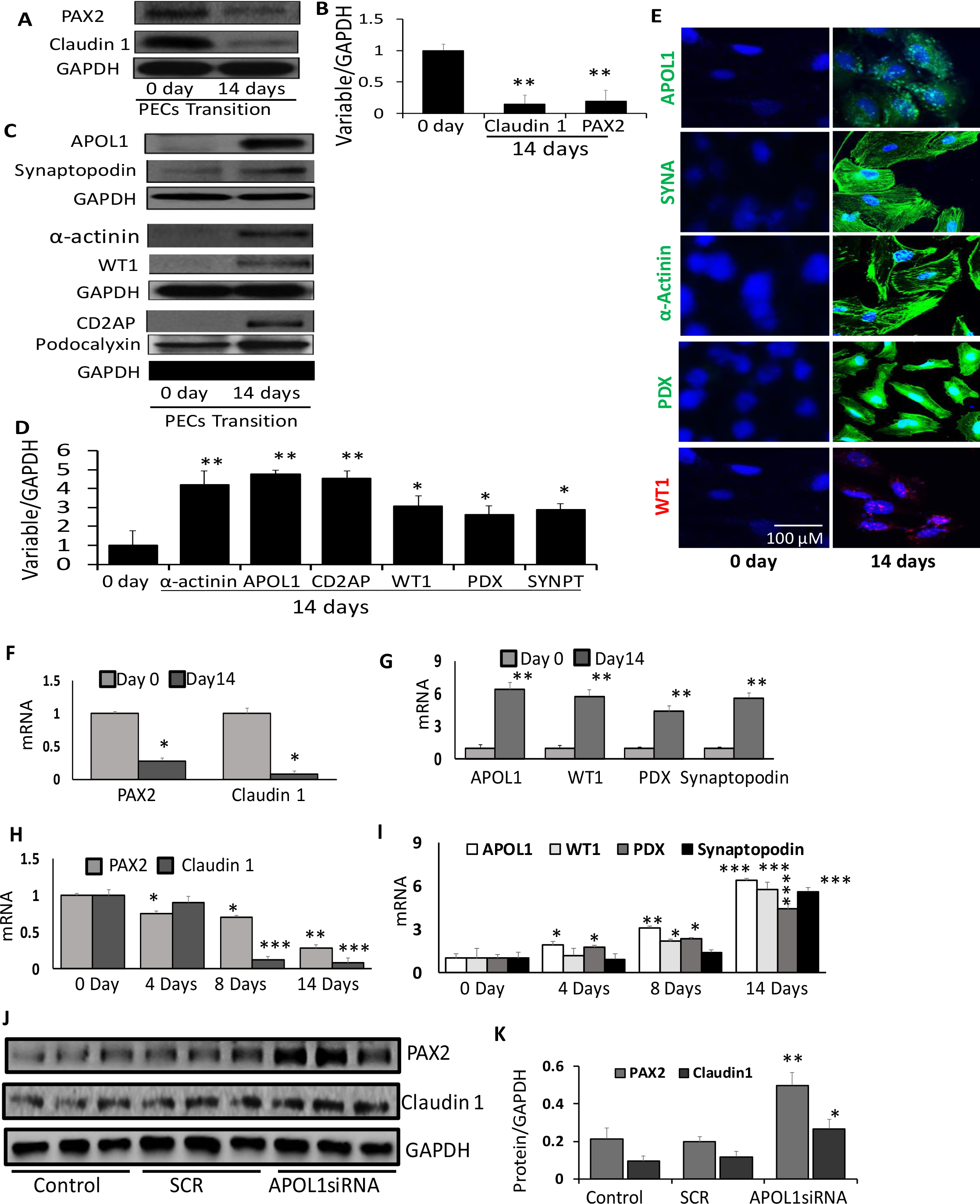
Absence or Presence of APOL1 Determines PECs Molecular profile. A. PECs were incubated in media for 14 days at 37°C (n=3). Protein blots of control (0 day) and transited (Tr)-PECs (14 days) were probed for PAX2 and reprobed for Claudin 1 and GAPDH. Representative gels are displayed. B. Cumulative densitometric data from protein blots from 2A are displayed in a bar diagram. *P<0.05 compared with respective 0 day; **P<0.01 compared to respective 0 day C. Protein blots from cellular lysates prepared in protocol A, were probed for APOL1 and re-probed for synaptopodin and GAPDH (n=3). Protein blots from the same lysates were probed for α-Actinin and re-probed for WT1 and GAPDH (n=3). Protein blots were also probed for CD2AP and re-probed for podocalyxin and GAPDH (n=3). Representative gels are displayed. D. Cumulative densitometric data of protein blots generated in 2C are shown in a bar diagram. **P<0.01 compared to respective 0 day. E. PECs grown on coverslips were fixed on 0 and 14 days and labeled for APOL1, synaptopodin, α-Actinin, podocalyxin (PDX) and WT1. Subsequently, PECs were examined under a confocal microscope. Representative microfluorographs are displayed. F. RNAs were extracted from the lysates of protocol 2A. cDNAs were amplified with specific primers for PAX2 and Claudin 1. *P<0.01 compared to respective 0 day. G. cDNAs from the 2F (above, RNAs) were amplified with specific primers for APOL1, WT1, podocalyxin (PDX), and synaptopodin. **P<0.01 compared with respective 0 day. H. PECs were incubated in media for variable time periods (0, 4, 8, and 14 days) at 37^°^C. RNAs were extracted and cDNAs were amplified with specific primers for PAX2 and Claudin 1. *P<0.05 compared to respective 0 day. **P<0.01 compared with respective 0, 4, and 8 days; ***P<0.001 compared with respective 0, and 4 days. I. cDNAs obtained from the 2H (RNAS) were amplified with specific primers for APOL1, WT1, podocalyxin (PDX) and synaptopodin. *P<0.05 compared with respective 0 day. **P<0.05 compared with 0 and 4 days; ***P<0.01 compared with respective 0, 4, and 8 days. J. PECs were transfected with either scrambled (SCR) or APOL1 siRNAs (n=3). Protein blots control cells and transfected cells were probed for PAX2, Claudin 1, and GAPDH. Gels from three different lysate preparations are displayed. K. Cumulative densitometric data from the Figure 2I are shown as a bar diagram. *P<0.05 compared with respective control and SCR; **P<0.01 compared with respective control and SCR.

To determine the role of APOL1’s (mRNA) absence in PECs’ phenotype, we evaluated the effect of the silencing *APOL1* mRNA on the expression of PEC markers. PECs were transfected with either scrambled (SCR) or APOL1 siRNAs (n=3). Protein blots probed for PEC markers from three different lysates are displayed in Fig. 2J. Cumulative data are shown in a bar diagram (Fig. 2K) APOL1 (mRNA)-silenced PECs displayed accentuated expression of PEC markers. These findings suggest that lack of APOL1 favors the PEC molecular phenotype profile. Further, it confirms our notion that functional APOL1 protein is present in PECs but remains undetectable with the use of current tools.

We examined whether HIV and IFN-y (known stimulators of APOL1; 4, 20) had a potential to induce APOL1 expression in PECs. PECs were treated with IFN-γ or transduced with HIV (NL4-3) (n=3). RNAs were assessed for miR193a and *APOL1*. IFN-γ(Fig. 3A, Lower panel) and HIV (Fig. 3B, Lower panel), both down regulated miR193a expression but up regulated *APOL1* mRNA expression (Figs. AE and 3B, Upper panels).

**Fig.3.**
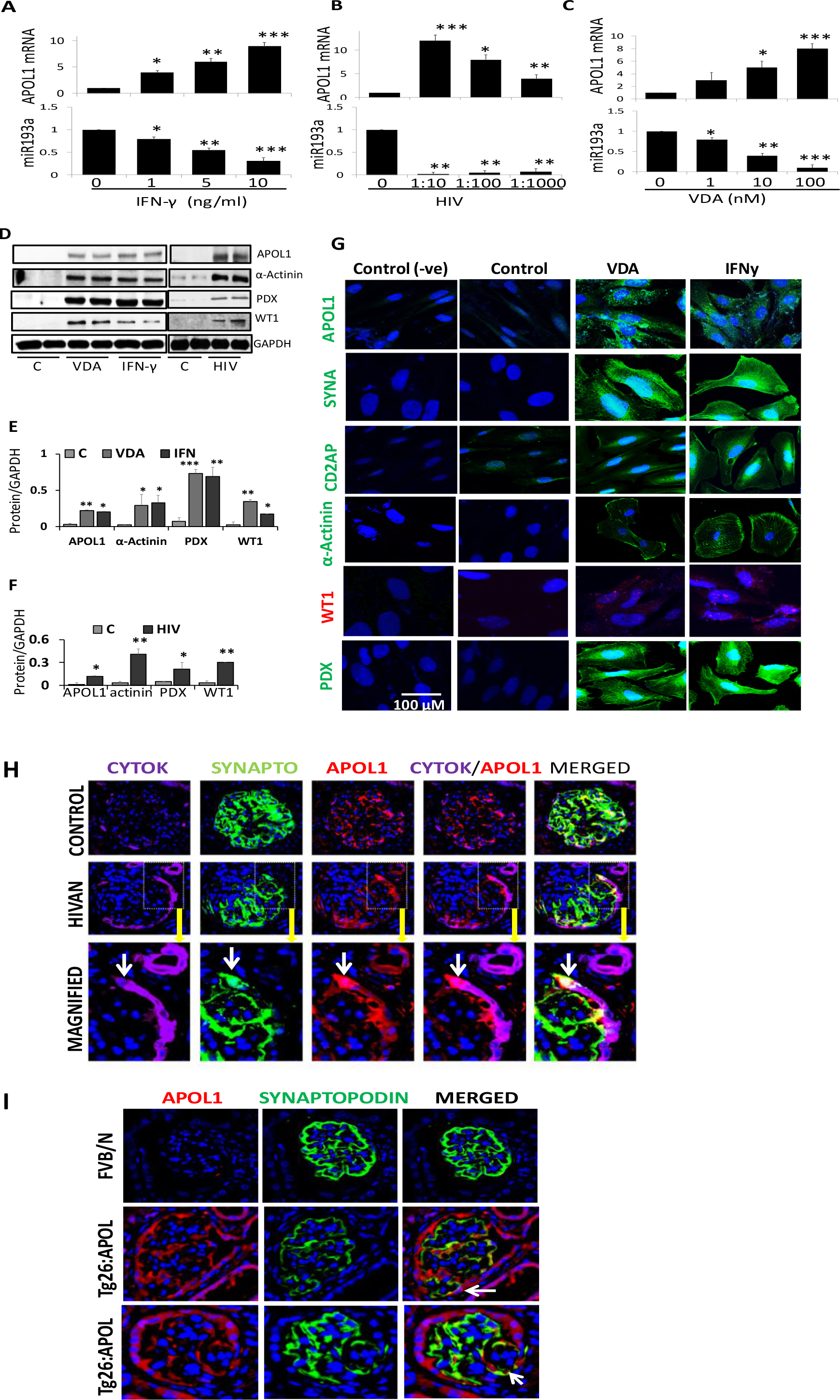
Role of APOL1 in PECs Transition. A. To determine the dose response effect of IFN-γ on miR193a and transcription of APOL1, PECs were incubated in media containing different concentration of IFN-γ (0, 1, 5, and 10 ng/ml) for 48 hours and assayed for miR193a (n=3). cDNAs were amplified with a primer specific for APOL1. Cumulative data on the expression of miR193a and APOL1 mRNA are shown in the lower and the upper panels, respectively. *P<0.05 compared with respective IFN, 0 ng/ml; **P<0.01 compared with respective IFN, 1.0 ng/ml; ***P<0.001 compared with respective IFN, 0 ng/ml. B. To determine the effect of HIV on miR193a and APOL1 expression, PECs were transduced with different concentrations of HIV (NL4-3) (n =3). After 48 hours, RNAs were assayed for miR193a. cDNAs were amplified for APOL1. Cumulative data on the expression of miR193a and APOL1 mRNA are shown in the lower and the upper panel, respectively. *P<0.05 compared with respective HIV, 0 and 1:10; **P<0.01 compared with respective HIV, 0; 1:10; ***P<0.001 compared with HIV, 0. C. To determine the effect of VDA on miR193a and APOL1 expression, PECs were incubated in media containing different concentrations of VDA (EB1089, 0, 1, 10, and 100 nM; 0 containing only vehicle) for 48 hours (n=3). RNAs were assayed for miR193a. cDNAs were amplified for APOL1. Cumulative data on the expression of miR193a and APOL1 mRNA are shown in the lower and the upper panel, respectively. *P<0.05 compared with respective VDA, 0 and 100 nM; **P<0.01 compared with respective VDA, 0, 1.0, and 100 nM, ***P<0.001 compared with respective 0. D. To determine the effect of APOL1 induction in PECs, PECs were incubated in media containing either vehicle (control), VDA (100 nM), or IFN-γ (10 ng/ml) for 48 hours (n=3). In parallel sets of experiments, PECs were transduced with vector (C) or HIV (n=3). Protein blots were probed for APOL1, WT1, podocalyxin, α-Actinin, and reprobed for GAPDH. Gels from two different lysates are shown. E. Cumulative densitometric data from the cells treated with VDA and IFN-γ in the protocol H. *P<0.05 compared to respective control; **P<0.01 compared to respective control; ***P<0.001 compared to respective control. F. Cumulative densitometric data from the cells transduced with vector (C) and HIV. *P<0.05 compared with respective control; **P<0.01 compared with respective control. G. PECs grown on coverslips were treated under similar conditions (as of 3D) and labeled for PECs’ transition markers. Representative microfluorographs are displayed. H. Paraffin fixed renal biopsy specimens of control and HIVAN patients colabeled for cytokeratin, synaptopodin, and APOL1. Representative fluoromicrographs are displayed. A glomerulus in a control patient did not show any expression of APOL1 by PECs but a glomerulus in a HIVAN patient displayed APOL1 expression by PECs. An occasional PEC also displayed colabeling for APOL1 and synaptopodin (indicated by white arrows). Mag. X200. Magnified inset X 800 I. Renal cortical sections of control (FVB/N) and HIV transgenic mice expressing APOL1 (Tg26: APOL1) were co-labeled for APOL1 and synaptopodin (n=4). Representative microfluorographs are shown. Foci of PECs displayed expression of synaptopodin (indicated by white arrows). Mag. X150

Since vitamin D has also been shown to down regulate miR193a expression in PECs (17), we evaluated the effect of a VDR agonist on APOL1 induction in PECs. PECs were treated with VDA and RNAs were evaluated for miR193a and *APOL1*. VDA not only down regulated miR193a expression (Fig. 3C, Lower panel) but also enhanced *APOL1* expression (Fig. 3C, Upper Panel).

To examine whether APOL1 inducers would also initiate PECs transition, PECs were incubated in media with either vehicle (control, DMSO), VDA, or IFN-γ. In parallel sets experiments (n=3), PECs were also transduced with either vector (C) or HIV. Protein blots probed for APOL1 and reprobed for transition markers from two different lysates are displayed in Fig. 3D. Cumulative data are shown in a bar diagram (Figs. 3E and 3F). PECs grown on coverslips were treated similarly and immunolabeled for different transition proteins. Representative fluoromicrographs are shown in Fig. 3G and Sup. Fig. 1. IFN-γ, VDA, and HIV not only induced the expression of APOL1 but also stimulated the expression of PECs’ transition markers.

To determine whether HIV induces APOL1 in PECs in patients with HIV-associated nephropathy (HIVAN), we carried out co-labeling for cytokeratin (PEC marker), synaptopodin (PD marker) and APOL1 in renal biopsy specimens of control and HIVAN patients. Representative fluoromicrographs are displayed in Fig. 3H. Glomeruli in control patients did not show any expression of APOL1 by PECs. However, Glomeruli in HIVAN patients displayed APOL1 expression by PECs. An occasional PEC also displayed co-labeling for APOL1 and synaptopodin suggesting transition of PEC.

To confirm the role of APOL1 in PECs’ transition *in vivo*, renal cortical sections of HIV transgenic mice expressing APOL1 (Tg26: APOL1) and FVB/N (control) mice were co-labeled for APOL1 and synaptopodin (n=4). Representative microfluorographs are shown in Fig. 3I. Foci of PECs displayed expression of synaptopodin. These findings suggest that APOL1 carry potential of stimulating PECs’ transition *in vivo*.

The question arises if APOL1 is an inducer of transition markers, it could do so in other kidney cells (which do not express APOL1) as well. To test our hypothesis, HEKs were transfected with either empty vector (EV) or APOL1 plasmids (n=3). Protein blots/cDNAS were assayed for APOL1 and transition markers. Representative gels from two different lysates are displayed in Fig. 4A. Cumulative data (protein blots in in Fig. 4B and mRNAs in Figs. 4C and Fig. 4D) are shown in bar diagrams. APOL1 induction decreased PEC marker, however, increased PECs’s transition markers in HEKs.

**Fig. 4.**
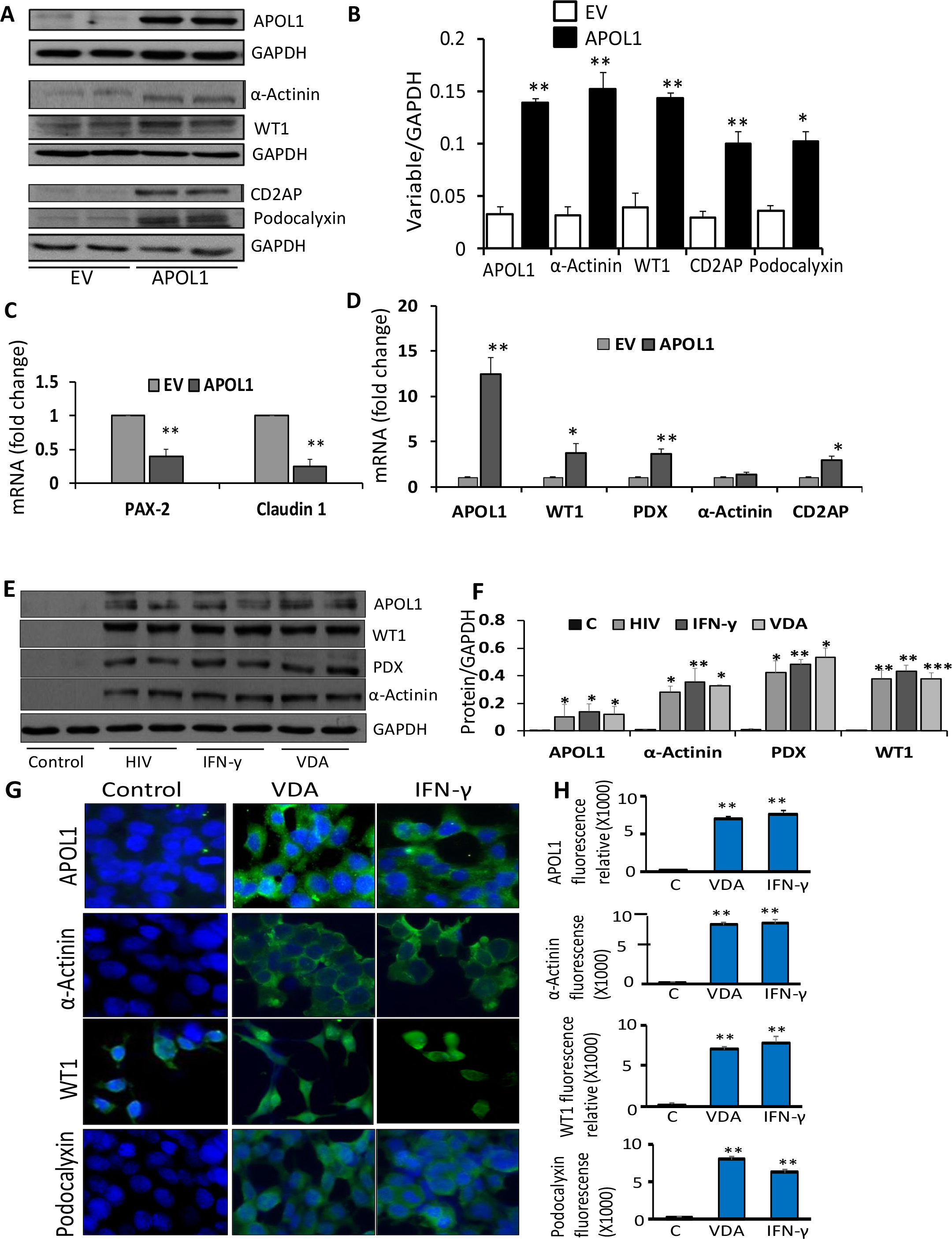
APOL1 induces transition markers in cells lacking APOL1. A. To determine the effect of APOL1 induction on transition markers, HEKs were transfected with empty vector (EV) or APOL1 plasmids (n=3). Protein blots were probed for APOL1 and reprobed for GAPDH. The same cellular lysates were probed for α-Actinin, WT1, CD2AP, podocalyxin and reprobed for GAPDH. Gels from two different lysates are displayed. B. Cumulative densitometric data from the protocol 3A, are displayed in a bar diagram.*P<0.05 compared with respective EV; **P<0.01 compared with respective EV. C. RNAs were extracted from the lysates of the protocol A and cDNAs were amplified for PAX2 and Claudin 1. Cumulative data are shown in a bar diagram. **P<0.01 compared with respective EV. D. RNAs were extracted from the lysates of the protocol A and cDNAs were amplified for APOL1, WT1, podocalyxin (PDX), α-Actinin, and CD2AP. Cumulative are shown in a bar diagram. *P<0.01 compared to respective EV; **P<0.01 compared to respective EV. E. To determine the effect of APOL1 induction on transition markers, HEKs were either transduced with HIV (NL4-3) or treated with IFN-γ (10 ng/ml) or VDA (100 nM) for 48 hours (n=3). Protein blots were probed for APOL1, WT1, podocalyxin, α-Actinin, and reprobed for GAPDH. Gels from two different lysates are shown. F. Cumulative densitometric data from the protein blots of the protocol 4E are shown in a bar diagram. *P<0.05 compared with respective control; **P<0.01 compared with respective control; ***P<0.001 compared with respective control. G. HEKs grown on coverslips were either transduced with HIV (NL4-3) or treated with IFN-γ (10 ng/ml) or VDA (100 nM) for 48 hours (n=3). Subsequently cells were labeled for APOL1, podocalyxin (PDX), α-Actinin, and WT1. Representative microfluorographs are displayed. H. Quantification of fluorescence data (25 - 40 cells in each group) from 4G are shown in bar graphs. **P<0.01 compared to respective control. I. RNAs were extracted from the lysates of the protocol 4E. cDNA were amplified with specific primers for APOL1, WT1, α-Actinin, and podocalyxin (PDX). Cumulative data (n=3) are shown in bar graphs. *P<0.05 compared with respective control. J. RNAs were extracted from the lysates of the protocol 4E. cDNA were amplified with specific primers for PAX2 and Caludin 1. Cumulative data (n=3) are shown in a bar diagram. *<0.05 compared to respective control; **P<0.01 compared to respective control.

To test whether APOL1 inducers could also induce APOL1 and associated downstream effects in cells lacking APOL1, HEKs were incubated in media containing vehicle (C, DMSO), IFN-γ, or VDA; in parallel sets of experiments (n=3), HEKs were transduced with HIV. Proteins and RNAs were extracted. Blots probed for APOL1 and reprobed for transition markers from two different lysates are displayed in Fig. 4E. Cumulative data are shown in a bar diagram (Fig. 4F). PECs grown on coverslips were treated under similar conditions. Subsequently, cells were labeled for transition makers. Representative fluoromicrographs are displayed in Fig. 4G. Fluorescence quantification data are shown in Fig. 4H. cDNAs were amplified for PECs’ and transition makers and data are shown in bar diagrams (Fig. 4I and 4J). HIV, IFN-γ, and VDA not only induced the expression of APOL1 but also stimulated transcription and translation of PECs’ transition markers in HEKs. Additional, data (FACS analysis) on VDA induced PEC transition markers in HEKs are shown in Sup. Fig. 2.

## APOL1 is critical for functionality of APOL1-miR193a axis

To test whether APOL1 plays a critical role in PECs’ transition, Tr-PECs were transfected with scrambled (SCR) or APOL1 siRNA. Control and experimental cells were treated with either vehicle (C), VDA, IFN-γ, miR193a inhibitor or transduced with HIV (n=4). Protein blots were probed for APOL1 and reprobed for transition markers. Representative Western blots are displayed in Figs. 5A and 5C. Cumulative data are shown in bar diagrams (Figs. 5B and 5D). VDA, IFN-γ, and miR193a inhibitor enhanced the expression of APOL1 and transition markers. However, the silencing of APOL1 down regulated the expression of transition makers despite inhibition of miR193a.

**Fig.5.**
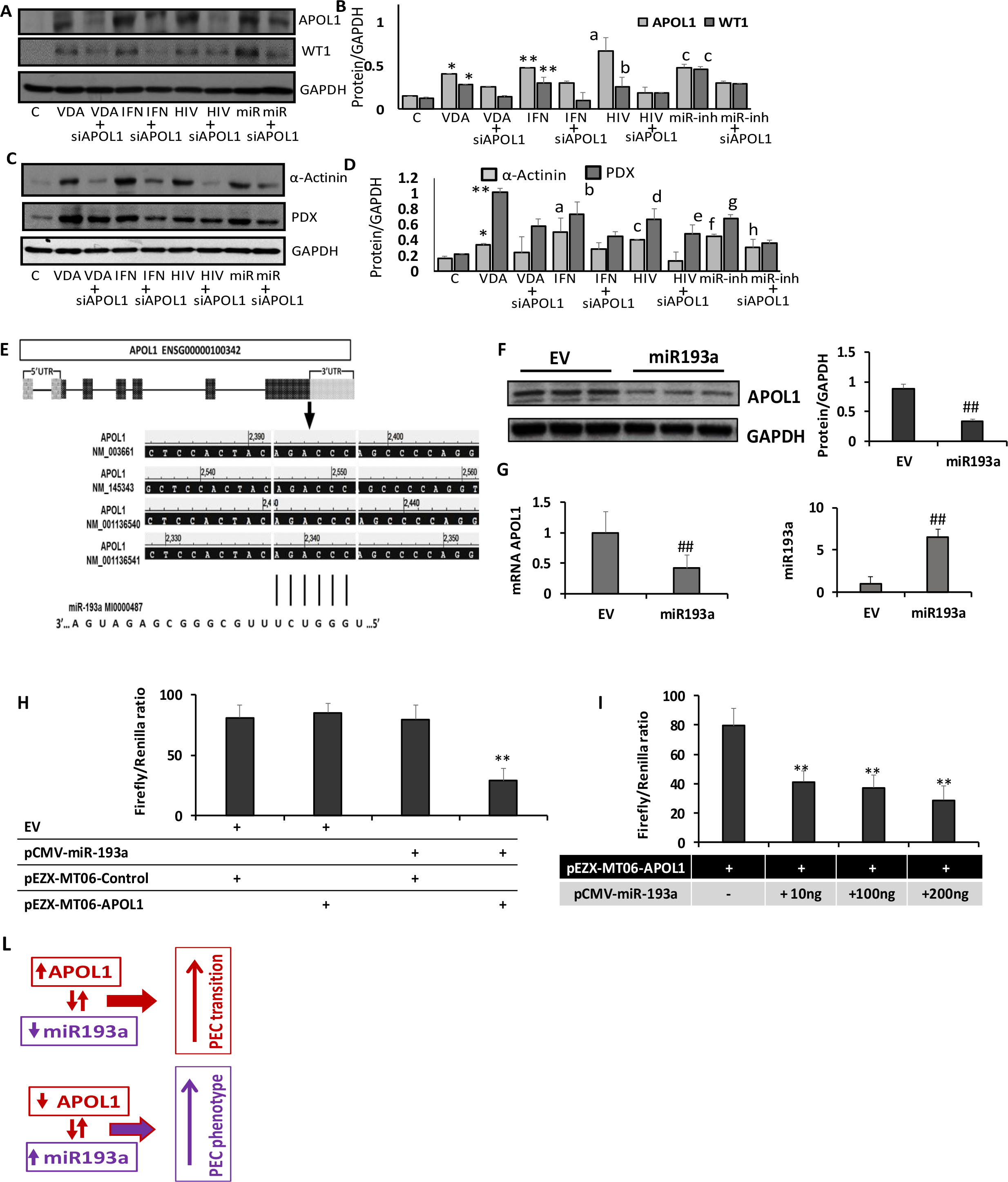
APOL1 is critical for PECs’ transition. A. To determine whether APOL1 is critical for transition in human PECs, Tr-PECs were transfected with scrambled (SCR) or APOL1 siRNA. Control and transfected cells were incubated in media containing either vehicle, VDA (10 nM), IFN-γ (10 ng/ml), or miR193a inhibitor (miR, 25 nM) (n=4). In parallel sets of experiments, control and transfected cells were transduced with HIV (NL4-3) (n=4). Protein blots were probed for APOL1 and reprobed for WT1, and GAPDH. Representative blots are displayed. B. Cumulative densitometric data of protein blots from the 4A are shown in bar graphs. *P<0.05 compared with respective C and VDA +siAPOL1; **P<0.05 compared with respective C and IFN + siAPOL1; ^a^P<0.01 compared with respective C and HIV + siAPOL1; ^b^P<0.05 compared with respective C; ^c^P<0.01 compared with respective C and miR + siAPOL1. C. Protein blots from the cellular lysates used in the protocol 4A, were probed for α-Actinin, podocalyxin (PDX) and GAPDH (n=4). Representative blots are displayed. D. Cumulative densitometric data of protein blots from the protocol 4C are shown in a bar diagram. *<0.05 compared with respective C, **P<0.01 compared with respective C and VDA +siAPOL1; ^a^P<0.05 compared with respective C and IFNγ + siAPOL1; ^b^P<0.01 compared with respective C and IFN + siAPOL1; ^c^P<0.05 compared with respective C and HIV + siAPOL1; ^d^P<0.01 compared with respective C; ^e^P<0.05 compared with respective C; ^f^P<0.05 compared with respective C and miR +siAPOL1; ^g^P<0.01 compared with respective C and miR193a + siAPOL1; ^h^P<0.05 compared with respective C. E. In silico computational algorithms based schematic representation of predicted miR193a target site in APOL1 3’UTR. F HepG2 cells were transiently transfected with either empty vector of miR293a plasmids. Protein blots probed for APOL and GAPDH. Gels from there different lysates are displayed. Cumulative data are shown in bar graphs. ^##^P<0.01 vs. EV. G. RNAs were extracted from the lysates of 5F. APOL1 mRNA and miR193a were analyzed using real time PCR normalized to GAPDH and U6 expression respectively. ##P<0.01 vs. respective EV. H. HEKs overexpressing pCMV-miR193a displayed decreased pEZX-MT06-APOL1 luciferase signal compared to EV. **P<0.01 vs. EV. I. Relative luciferase activity with different concentration of pCMV-miR193a. **p<0.01 vs. 0 concentration. L. Proposed schematic hypothesis of APOL1 dynamics in human PEC phenotype kinetics. APOL1 and miR193a display a reciprocally linked feedback loop. Up regulation of APOL1 and down regulation of miR193a feedback each other and initiate PECs transition; conversely, down regulation of APOL1 with up regulation of miR193a in a reciprocal manner assures PECs phenotype.

Our data suggest that miR193a is a potential target for APOL1. To validate our notion, complimentary APOL1 binding sites of miR193a were mapped by bioinformatics analysis (Fig. 5E). HepG2s were transiently transfected with either empty vector or miR193a plasmid and evaluated for APOL1 protein expression (Fig. 5F) and APOL1 mRNA/miR193a expression (Fig. 5G). HEKs overexpressing pCMV-miR193a displayed significantly decreased pEZX-MT06-APOL1 luciferase signal compared to empty vector (Fig. 5H), suggesting a putative interaction between miR-193a and APOL1. Relative luciferase activity with different concentrations of pCMV-miR-193a are shown in Fig. 5I.

PECs have been demonstrated to participate in PD homeostasis in mice, which do not carry APOL1 (21–23); nonetheless, the numbers of PECs capable of transiting to PDs shown to be limited and transited during juvenile period only (21–24). It is plausible that the majority of PECs in adult mice are not able to participate in PD homeostasis as a consequence of lack of APOL1. It would be worth studying this aspect in APOL1 transgenic mice in future studies.

In summary, both APOL1 and miR193a are reciprocally linked with a feedback loop in human PECs (Fig. 5L). Up regulation of miR193a sustains the PEC phenotype through down regulation of APOL1 and WT1, whereas; down regulation of miR193a initiates PECs' transition through up regulation of APOL1.

## Methods

### Human PECs, HEKs and HepG2s

Immortalized human PECs were initially obtained from Prof. Catherine Meyer-Schwesinger, University Medical Center, Hamburg-Eppendorf, Hamburg, Germany. PECs were grown in 1:1 EGM endothelial growth medium supplemented with 5% FCS, 0.4% bovine brain extract, 0.1% hEGF, 0.1% hydrocortisone, 0.1% gentamicin and amphotericin B, 100 units/ml penicillin, 100 mg/ml streptomycin (Lonza, Basel, Switzerland), and RPMI 1640 supplemented with 10% FCS and 1× ITS at 33°C (5% CO_2_ incubator). For transition (differentiation), PECs were cultured for 14 days in EGM endothelial growth medium and RPMI 1640 with the added supplement described above. During transition, PECs were seeded on laminin- or fibronectin-coated dishes (BD Biocoat Cellware; BD Biosciences), and the medium was supplemented with a special induction medium containing vitamin D_3_ (100 *μ*M; Sigma-Aldrich), retinoic acid (100 *μ*M; Sigma-Aldrich), and dexamethasone (0.1 μM; Sigma-Aldrich).

HEKs were purchased from ATCC and cultured in DMEM supplemented with 10% FCS and 25 mM Hepes. HepG2 cells were gifted by Prof. Sanjeev Gupta (Liver Research Center, Albert Einstein College Medicine, Bronx, NY). These cells were cultured in DMEM supplemented with 10% FCS and 25 mM Hepes.

### HIV(Tg26): APOL1 Transgenic mice

APOL1 transgenic mice were generated by cloning APOL1 cDNA into the vector (TetOn3G; http://www.clontech.com/US/Products/Inducible_Systems/Tetracycline-Inducible_Expression/Tet-On_3G). The transgene (TetON-APOL1) was injected into FVB/N oocyte (Cold Spring Harbor Laboratory, Cold Spring, NY). APOL1 +ve transgenic mice were identified by genomic PCR analysis using transgene specific primers. APOL1 transgenic mice were cross-bred with Tg26 (HIV transgenic, a gift from Prof. Paul Klotman, University Texas, Houston) mice. Tg26 mice are also on FVB/N background. Transgene expression was induced by doxycycline containing feed (20 mg/Kg, BioServe). Animal studies were approved by the Animal Care Committee of Feinstein Institute for Medical Research, Northwell Health, New York.

### Human Renal biopsy data

Archived biopsy specimens from the patients of HIV-associated nephropathy and control patients (renal transplant kidney donors) were used. Institute review board of Feinstein Institute for Medical Research, Northwell Health, exempted use of these archived tissues

### Immunofluorescence of PECs and HEKs

Cells were grown on fibronectin-coated coverslips. After differentiation or inductions, cells were washed with 1xPBS and fixed in 4% paraformaldehyde for 10 min at room temperature. Cells were washed three times with 1xPBS and permeabilized with 0.5% Triton X-100. After another washing step with 1xPBS, cells were incubated in 5% BSA (Bovine Serum Albumin) for 30 min at room temperature. Cells were incubated with the following primary antibodies: m-APOL1 (1:100, Proteintech), rb-PAX2 (1:200, Abcam), rb-Cladudin1 (1:200, Abcam), rb-WT1 (1:100, Abcam), rb-podoclyxin (1:200, Life technologies) and m-α-Actinin (1:200, Santa Cruz biotechnologies) for overnight at 4°C. After incubation, cells were washed three times with 1xPBS and stained with fluorochrome (AF488, 1:200) conjugated secondary antibody to goat anti-rabbit and donkey anti-mouse (Molecular Probes) for 30 min at room temperature and nuclear stain Hoechst 33342. For each set of experiment, negative control was prepared with the use of isotype IgG in place of primary antibody followed by secondary antibody. Staining was visualized under a confocal microscope.

### Immunofluorescence of kidney tissue

Human renal biopsy slides (control and HIVAN patients) and slides of renal cortical sections from FVB/N (control) and HIV: APOL1 transgenic (Tg26: APOL1) mice were placed into 100% xylene for 5 minutes for 4 times, then slides were placed into 100% ethanol for 5 minutes twice, then into 70% ethanol 5 minutes twice, then into 50% ethanol for 5 minutes twice. Slides were washed with ddH_2_O for 1 minute. Now slides were subjected into 1x retrieve-All antigen unmasking system buffer (Covance, Dedham) at 100 °C for 90 minutes. Slides were placed at room temperature for 20-30 minutes for cooling. Then washed with 1x PBS. Slides were subjected into 0.3% triton X-100 for 20 minutes at room temperature. Slides were blocked with 2% BSA for 2 hours. After blocking, primary antibody (cytokeratin at 1:50 dilution, synaptopodin at 1:50 dilution, and APOL1 at 1:100 dilution) were added for overnight at 4 °C. Next day, slides were washed with 0.1% triton X-100 for 5 minutes thrice on shaker at room temperature. Secondary antibody was added with fluorescence conjugated at 1:500 dilutions for 1 hour at room temperature. Nuclei were stained by DAPI. Slides were washed with 0.1% triton X-100 for 5 minutes thrice on shaker at room temperature and mounted for confocal microscopy.

### qPCR

Total RNA was isolated from hPECs, HEK, HepG2 and human podocytes with TRIzol reagent (Invitrogen, USA). cDNA was synthesized using 1 *μg* of total RNA treated with High-Capacity cDNA Reverse Transcription kit (Thermofisher, USA) according to the manufacturer's instructions. Real-Time PCR was performed using one-step iTaq^TM^ Universal Syber Green kit (BIO-RAD, USA) using specific primers obtained from Invitrogen Life Technologies. *GAPDH* fw 5' CCC ATC ACC ATC TTC CAG GAG '3; rev 5' GTT GTC ATG GAT GAC CTT GGC '3, *WT1* fw 5' CGAGAGCGATAACCACACAACG '3; rev 5' GTCTCAGATGCCGACCGTACAA '3, Synaptopodin fw 5' CTT CTC CGT GAG GCT AGT GC '3; rev 5' TGA GAA AGG CTT GAAA GG '3, *Podocalyxin* fw 5' TGT TTT GTT AGA TGA GTC CGT AGT A '3; rev 5' CGC TGC TAC TGT CA '3, *α-Actinin-4* fw 5' GTT CTC GAT CTG TGT GCC TG '3; rev 5' GAC CTG GAC CC '3, Claudin-1 fw 5' TGG GGG TGC GAT ATT TCT TC '3; rev 5' CCT CCC AGA AGG CAG AGA GA '3, *PAX2* fw 5' GGC TGT GTC AGC AAA ATC CTG '3; rev 5' TCC GGA TGA TTC TGT TGA TGG '3, *APOL1* fw 5' ATC TCA GCT GAA AGC GGT GAAC '3; rev 5' TGA CTT TGC CCC CTC ATG TAAG '3, *CD2AP* fw 5' AAG GGT GGC TGG AAG GAG AAC '3; rev 5' ATG CCT TTC CCG TTT GAT GG '3. For miR quantification, the total RNA was isolated with miRVana miRNA Isolation Kit and TaqMan-based detection primers miR-193a and U6 (internal control) (Thermofisher Scientific, USA) were used for Real-time amplification (ABI-7500, Applied Biosystems). Relative quantification of gene expression was calculated using the ΔΔCT method.

### Western Blot Analysis and Antibodies

Protein blots of control and experimental cells were performed as previously described (4). Briefly, protein lysates were prepared using RIPA lysis buffer (Millipore, USA). After boiling for 15 minutes, equal volume of cell lysates were electrophoresed using 10-12% of SDS-PAGE gels. Separated proteins were transferred to PVDF (EMD Millipore) membranes and processed further for immunostaining with primary antibodies against APOL1 (mouse monoclonal, ProteinTech), WT1 (rabbit polyclonal, SantaCruz), Synaptopodin (polyclonal rabbit, Santa Cruz), CD2AP (mouse monoclonal, SantaCruz), Podocalyxin (rabbit polyclonal, SantaCruz), α-Actinin (mouse monoclonal, Abcam), PAX2 (rabbit polyclonal, Abcam) and -Claudin-1 (rabbit polyclonal, Abcam). All the blots were incubated with primary antibodies incubated at 4°C overnight and subsequently with horseradish peroxidase-labeled appropriate secondary antibodies (1:3000, Santa Cruz). The blots were developed using a chemiluminescence detection kit (Pierce, Rockford, IL) and exposed to X-ray film (Eastman Kodak, Rochester, NY) or chemiluminescence reader and image lab software (Chemidoc MP, Bio-Rad). M-m-GAPDH (1:3000, Santa Cruz Biotechnology) was used as an internal control to confirm equal protein loading.

### Luciferase assay

Complimentary APOL1 binding sites of miR193a were mapped by bioinformatics analysis using three different miRNA databases (microrna.org; mirdb.org and TargetScan). The APOL1 wild-type 3'UTR (pEZX-MT06-APOL1,HmiT055026-MT06) and pEZX-MT06-Control (scrambled non-specific sequence 3'UTR control vectors, CmiT000001-MT06) containing firefly luciferase and *Renilla* luciferase as tracking genes were purchased from GeneCopoeia. Luc-Pair™ Duo-Luciferase HS Assay Kit was used to measure the relative luciferase activity as described in manufacturer’s protocols. Briefly, cells were transiently co-transfected by using Lipofectamine 2000 with wild-type or control reporter 3’-UTR plasmids and miR-193a (pCMV-miR-193a) or negative miR (control, AM17110) in combination. After 48 hours of co-transfection the firefly luciferase activities were measured using the Duo-Luciferase HS Assay (GeneCopoeia).The relative luciferase activity was calculated by normalizing to Renilla luciferase. Results represent cumulative values of three independent experiments, each done in triplicates.

### Statistical analyses

Statistical comparisons were performed with the program PRISM using the Mann–Whitney *U* test for nonparametric data and the unpaired *t* test for parametric data. A *P* value<0.05 was accepted as statistically significant.

## Acknowledgements

This work was supported by grants RO1DK 098074, RO1DK084910, RO1 DK083931 (PCS) from National Institutes of Health, Bethesda, MD.

---

**Sup. Fig. 3.**
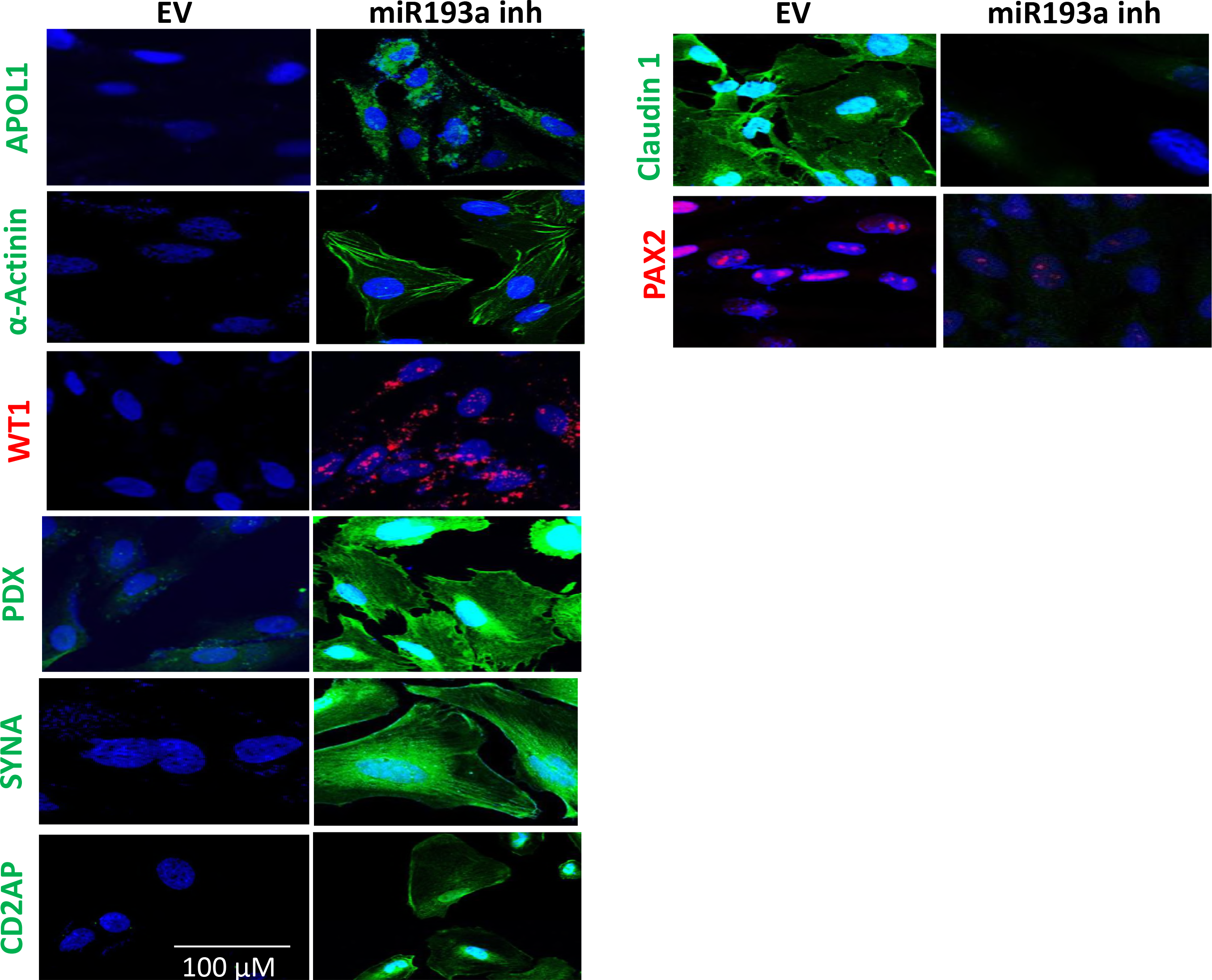
PECs grown on coverslips were transfected with either empty vector (EV) or miR193a inhibitor (inh) at 37°C. After 48 hours, PECs were labeled for the Indicated antibodies and examined under a confocal microscope. Representative microfluorographs are displayed.

## References

1. Vanhamme, L., F. Paturiaux-Hanocq, P. Poelvoorde, D.P. Nolan, L. Lins, J. Van Den Abbeele, A. Pays, P. Tebabi, H. Van Xong, A. Jacquet, N. Moguilevsky, M. Dieu, J.P. Kane, P. De Baetselier, R. Brasseur, and E. Pays. 2003. Apolipoprotein L-I is the trypanosome lytic factor of human serum. Nature 422: 83–87.

2. Kopp, J.B., G.W. Nelson, K. Sampath, R.C. Johnson, G. Genovese, P. An, D. Friedman, W. Briggs, R. Dart, S. Korbet, M.H. Mokrzycki P.L.Kimmel, S. Limou, T.S. Ahuja, J.S. Berns, J. Fryc, E.E. Simon, M/C. Smith, H. Trachtman, D. M. Michel, J. R. Schelling, D. Vlahov, M. Pollak, and C.A. Winkler. 2011. APOL1 genetic variants in focal segmental glomerulosclerosis and HIV-associated nephropathy. J Am Soc Nephrol 22: 2129–2137.

3. Friedman, D.J., J. Kozlitina, G. Genovese, P. Jog, and M.R. Pollak. 2011. Population-based risk assessment of APOL1 on renal disease. J Am Soc Nephrol22: 2098–2105.

4. Lan, X., A. Jhaveri, K. Cheng, H. Wen, M. A. Saleem, P.W. Mathieson, J. Mikulak, S. Aviram, A. Malhotra, K. Skorecki, and P.C. Singhal. 2014. APOL1 risk variants enhance podocyte necrosis through compromising lysosomal membrane permeability.Am J Physiol Renal Physiol. 307:F326–36.

5. Lan, X., H. Wen, R. Lederman, A. Malhotra, J. Mikulak, W. Popik, K. Skorecki K, and P.C. Singhal. 2015̤Protein domains of APOL1 and its risk variants.Exp Mol Pathol. 99:139–44.

6. Kruzel-Davila, E., R. Shemer, A. Ofir, I. Bavli-Kertselli, I. Darlyuk-Saadon, P. Oren-Giladi, W.G. Wasser, D. Magen, E. Zaknoun, M. Schuldiner, A. Salzberg, D. Kornitzer, Z. Marelja, M. Simons, and K. Skorecki K. 2016. APOL1-Mediated Cell Injury Involves Disruption of Conserved Trafficking Processes. J Am Soc Nephrol. 28:1117–1130.

7. Fu, Y., J.Y. Zhu, A. Richman, Y. Zhang, X. Xie, J.R. Das, J. Li, P.E. Ray, and Z. Han. 2016. APOL1-G1 in Nephrocytes Induces Hypertrophy and Accelerates Cell Death. J Am Soc Nephrol. 28:1106–1116.

8. Ma, L., J.W. Chou, J.A. Snipes, M.S. Bharadwaj, A.L. Craddock, D. D. Cheng, A. Weckerle, S. Petrovic, P.J. Hicks, A.K. Hemal, G.A. Hawkins, L.D. Miller, A.J. Molina, C.D. Langefeld, M. Murea, J.S. Parks, and B.I. Freedman. 2017. APOL1 Renal-Risk Variants Induce Mitochondrial Dysfunction. J Am Soc Nephrol. 28:1093–1105.

9. Khatua, A.K., A.M. Cheatham, E.D. Kruzel, P.C. Singhal, K. Skorecki, and W. Popik. 2015. Exon 4-encoded sequence is a major determinant of cytotoxicity of apolipoprotein L1. Am J Physiol Cell Physiol. 309:C22–37.

10. Olabisi, O.A., J.Y. Zhang, L. VerPlank, N. Zahler, S. DiBartolo 3rd, J.F. Heneghan, J.S. Schlöndorff, J.H. Suh, P. Yan, S.L. Alper, D.J. Friedman, and M.R, Pollak. 2016. APOL1 kidney disease risk variants cause cytotoxicity by depleting cellular potassium and inducing stress-activated protein kinases. Proc Natl Acad Sci U S A. 113:830–7.

11. Beckerman, P., J. Bi-Karchin, A.S. Park, C. Qiu, P.D. Dummer, I. Soomro, C.M. Boustany-Kari, S. S. Pullen, J.H. Miner, C̤ Hu, T. Rohacs, K. Inoue, S. Ishibe, MA Saleem, M.B. Palmer, A. M. Cuervo, J.B. Kopp, and K. Susztak. 2017.Transgenic expression of human APOL1 risk variants in podocytes induces kidney disease in mice. Nat Med. 23: 429–438.

12. Hayek, S.S., K.H.E. Hahm, V. Peev, M. Tracy, N.J. Tardi, V. Gupta, M.M. Altintas, G. Garborcauskas, N. Stojanovic, C.A. Winkle, M.S. Lipkowitz, A. Tin, L.A. Inker, A.S. Levey, M. Zeier, B.I. Freedman, J.B. Kopp, K. Skorecki, J. Coresh, A.A. Quyyumi, S. Sever, and J. Reiser. 2017. A tripartite complex of suPAR, APOL1 risk variants and α integrin on podocytes mediates chronic kidney disease.Nat Med. 23:945–953.

13. O'Toole, J.F., W. Schilling, D. Kunze, S.M. Madhavan, M. Konieczkowski, Y. Gu, L. Luo, Z. Wu, L.A. Bruggeman, and J.R. Sedor. 2017. ApoL1 Overexpression Drives Variant-Independent Cytotoxicity. J Am Soc Nephrol. doi: 10.1681/ASN.2016121322

14. Lazzeri, E. and P. Romagnani. 2015̤Podocyte biology: Differentiation of parietal epithelial cells into podocytes.Nat Rev Nephrol. 11:7–8.

15. Shankland, S.J., B. Smeets, J.W. Pippin, and M.J. Moeller.2014. The emergence of the glomerular parietal epithelial cell.Nat Rev Nephrol. 10:158–73.

16. Eng, D.G., M.W. Sunseri, N.V. Kaverina, S.S. Roeder, J.W. Pippin, and S.J. Shankland. 2015.. Glomerular parietal epithelial cells contribute to adult podocyte regeneration in experimental focal segmental glomerulosclerosis. Kidney Int. 88:999–1012.

17. Kietzmann, L., S.S. Guhr, T.N. Meyer, L. Ni, M. Sachs, U. Panzer, R.A. Stahl, M.A. Saleem, D. Kerjaschki, C.A. Gebeshuber, and C. Meyer-Schwesinger C. 2015. MicroRNA-193a Regulates the Transdifferentiation of Human Parietal Epithelial Cells toward a Podocyte Phenotype. J Am Soc Nephrol. 26:1389–401.

18. Mishra, A., K. Ayasolla, V. Kumal, X̤ Lan, H. Vashistha, R. Aslam, A. Hussain, S. Chowdhary, S. M. Shoshtari, N. Paliwal, W. Popik, M. A. Saleem, A. Malhotra,L.G. Meggs, K. Skorecki, and P. C. Singhal. 2018. Modulation of APOL1-miR193a Axis Prevents Podocyte Dedifferentiation in High Glucose Milieu. Am J. Physiol. Renal. In press.

19. Madhavan, S.M., J.F. O'Toole, M. Konieczkowski, S. Ganesan, L.A. Bruggeman, and JR Sedor. 2011. APOL1 localization in normal kidney and nondiabetic kidney disease. J Am Soc Nephrol. 22:2119–28.

20. Mikulak, J., F. Oriolo, F. Portale, P. Tentorio, X. Lan, M.A. Saleem, K. Skorecki, P.C. Singhal, and D. Mavilio. 2016. .Impact of APOL1 polymorphism and IL-1β priming in the entry and persistence of HIV-1 in human podocytes. Retrovirology. 13:63

21. Appel, D., D.B. Kershaw, B. Smeets, G. Yuan, A. Fuss, B. Frye, M. Elger, W. Kriz, J. Floege, and M.J. Moeller. 2009. Recruitment of podocytes from glomerular parietal epithelial cells. J Am Soc Nephrol. 20:333–43.

22. Berger, K., K. Schulte, P. Boor, C. Kuppe, T.H. van Kuppevelt, J. Floege, B. Smeets, and M.J. Moelle. 2014. The regenerative potential of parietal epithelial cells in adult mice.J Am Soc Nephrol. 25:693–70.

23. Berger, K. and M.J. 2014. Moeller MJ.Podocytopenia, parietal epithelial cells and glomerulosclerosis.Nephrol Dial Transplant. 29:948–5.

24. Shankland, S.J., B.S. Freedman and J.W. Pippin. 2017. Can podocytes be regenerated in adults? Curr Opin Nephrol Hypertens. 26:154–164, 2017

